# ASD-Associated *CNTNAP2* Variants Disrupt Neuronal Arborization Through Impaired Regulation by Ectodomain Shedding

**DOI:** 10.1101/2024.11.04.621898

**Authors:** Miguel Lobete, Leonardo E. Dionisio, Emmarose McCoig, Nicolas H. Piguel, Benjamin P. Spielman, Silvia Socas, Marc dos Santos, Cristina Boers-Escuder, Peter Penzes, M. Dolores Martin-de-Saavedra

## Abstract

Ectodomain shedding (ES) is a process by which a protease cleaves the extracellular portion of membrane-bound proteins, releasing soluble fragments that influence diverse cellular functions. ES is critical in neurodevelopment, plasticity, and neurodegenerative disorders, such as Alzheimer’s disease, and has recently been implicated in neurodevelopmental conditions, including autism spectrum disorders (ASD). Contactin-associated protein-like 2 (CNTNAP2) is an adhesion molecule regulated by ES, releasing a soluble ectodomain (sCNTNAP2) that enhances neuronal synchrony. CNTNAP2 is implicated in ASD, schizophrenia, and cortical dysplasia focal epilepsy syndrome (CDFE) and it is known to regulate neuronal arborization, as well as dendritic spine maturation and maintenance. However, little is known about how neuroplasticity impacts ES or the role of CNTNAP2 ES in dendritic arborization. Here, we show that the brain sheddome is enriched in shed ectodomains that regulate neuronal projections, and that its molecular and functional composition is modulated by sensory deprivation in a sex dependent manner, with a decrease in sCNTNAP2 levels observed only in male mice. Furthermore, we demonstrate that sCNTNAP2 promotes dendritic arborization, while ASD-associated CNTNAP2 variants present reduced sCNTNAP2 levels in culture and decreased neuronal branching. Together, these findings underscore the role of ES in neuroplasticity and ASD and reveal how CNTNAP2 genetic variations disrupt its regulation by ES, leading to altered dendritic branching.

## INTRODUCTION

Ectodomain shedding (ES) is a post-translational modification in which proteases cleave the extracellular domain of membrane-anchored proteins, releasing a soluble fragment. Although initially seen as a way to terminate protein function (1), recent studies have revealed additional roles for ES and its cleaved products, including the regulation of neurodevelopment (2, 3), plasticity (4, 5), and intercellular signaling (6, 7). Traditionally, ES studies focused on individual proteins; however, recent efforts have been placed on deciphering the molecular and functional composition of the “sheddome,” the group of proteins that are released by ES in specific tissues or samples. This shift has revealed that the neuronal, cortical, and cerebrospinal fluid (CSF) sheddomes might influence critical aspects of nervous system development, such as neuronal projection, axon guidance, axonogenesis, synapse assembly, and cell adhesion (7–9).

Emerging research also links ES to the pathophysiology of various conditions, ranging from cardiovascular (10) to neurodegenerative disorders (11–13). For instance, amyloidogenic processing of amyloid precursor protein (APP) through ES is a key event in Alzheimer’s disease (AD) pathology (14, 15), and ES regulates additional AD-associated risk factors, such as TREM2 (16, 17) and SORLA (18, 19). Given the sheddome’s involvement in neurodevelopmental processes, ES may also play a role in neurodevelopmental disorders, including autism spectrum disorders (ASD) and schizophrenia, which has been previously hypothesized (7), though this remains to be fully elucidated.

Neurodevelopment encompasses a series of highly coordinated processes— neurogenesis, cell migration, axonal and dendritic formation, synaptogenesis, synaptic refinement, and myelination—all shaped by sensory input (20–23). Yet, how sensory input might regulate ES and influence neurodevelopment-related cellular events remains largely unexplored.

CNTNAP2 (or Caspr2) is a type I transmembrane cell adhesion molecule of the neurexin superfamily, known to undergo ES and release a soluble ectodomain (sCNTNAP2) (7, 24). *CNTNAP2* has been implicated in many neurodevelopmental conditions including ASD, intellectual disability, language impairment, schizophrenia, and cortical dysplasia focal epilepsy (CDFE) (for review see references (25, 26)). Disruptions in *CNTNAP2* in ASD include translocations, inversions, insertions, and point mutations (27–31). Originally identified for clustering Kv1 channels in myelinated axons via interaction with Contactin 2 (32), CNTNAP2 also plays a role in neuronal arborization (33–35), spine maturation and maintenance (36, 37), and AMPA receptor regulation (36, 38). Notably, sCNTNAP2 modulates neuronal synchrony in cultured neurons and in brain slices (7), and its levels are decreased in the cerebrospinal fluid of individuals with ASD (39). Since alterations in neuronal synchrony are a hallmark of ASD and other neurodevelopmental disorders (40–42), these findings suggest a potential link between CNTNAP2 ES and the disruptions in neuronal communication and behavior observed in these conditions. However, it remains unclear how CNTNAP2’s ES regulates dendritic arborization and whether ASD-linked genetic variations in CNTNAP2 affect this function.

This study investigates the role of ES in neurodevelopmental conditions by examining the impact of sensory deprivation on the molecular and functional composition of the sheddome and by exploring how ASD-associated CNTNAP2 variants influence ES regulation and dendritic arborization, offering new insights into CNTNAP2’s role in neurodevelopment and ASD pathophysiology.

## RESULTS

### The cortical sheddome is enriched in proteins regulating neuronal projections and ASD risk factors

Since the sheddome is enriched in proteins essential for neurodevelopment, we analyzed the cortical soluble fraction extracted from 15-day-old male and female mice (3 animals per sex)—a developmental stage marked by active neuroplasticity and probably increased ES. In contrast, most previous sheddome studies have focused on adult male brain tissue or adult cerebrospinal fluid (CSF) (7–9). We processed the cortices according to our published protocol (8). Briefly, cortices were dissected, homogenized using a Dounce homogenizer in a detergent-free buffer, then centrifuged at low speed to remove big organelles, nuclei, and cell debris. The resulting supernatants were ultracentrifuged for 2h at 100,000 g for 2 h at 4°C. The soluble fractions were analyzed by Liquid Chromatography-Tandem Mass Spectrometry (LC-MS/MS), with the sheddome isolated by bioinformatically selecting those proteins containing transmembrane domains or glycosylphosphatidylinositol (GPI) anchors according to the UniProt database. Out of the 4432 proteins detected in the soluble fraction, 274 contained a transmembrane domain or a GPI anchor and were present in at least 3 of the six samples, comprising the soluble fraction sheddome (**Fig. 1A**). Gene ontology (GO) analysis of the 274 proteins in the sheddome revealed several enriched biological processes related to neurodevelopment. These processes included cell migration, synapse assembly, organization and maturation. Notably, specific processes associated with neuronal projection development and extension, including processes specific to axons (axonogenesis and axon guidance) and dendrite morphogenesis (**Fig. 1B**). Moreover, the term ‘cell adhesion’ exhibited the highest adjusted *P*-value (**Fig. 1B**), reinforcing the notion that cell adhesion proteins are among those that undergo ES. We then performed a disease enrichment analysis on the cortical sheddome. Interestingly, we observed a significant enrichment of ASD risk-gene encoded proteins cataloged by the Simons Foundation Autism Research Initiative (SFARI). We corroborated that enrichment in ASD-associated *de novo* variation genes (43) and those identified in a GWAS meta-analysis of individuals with autism (44). Similarly, schizophrenia (SZ) risk-gene-encoded proteins (43) were also enriched in the cortical sheddome. Finally, we built a protein-protein interaction (PPI) network in which we highlighted proteins involved in cell adhesion (blue), neuron projection development (green), and CNS development (red) based on our GO analysis. The depicted PPI networks include many proteins that are known to undergo ES, such as APP, APLP1 and 2, L1CAM, various members of the CADM family, as well as neurexins and neuroligins (**Fig. 1D**).

**Figure 1.**
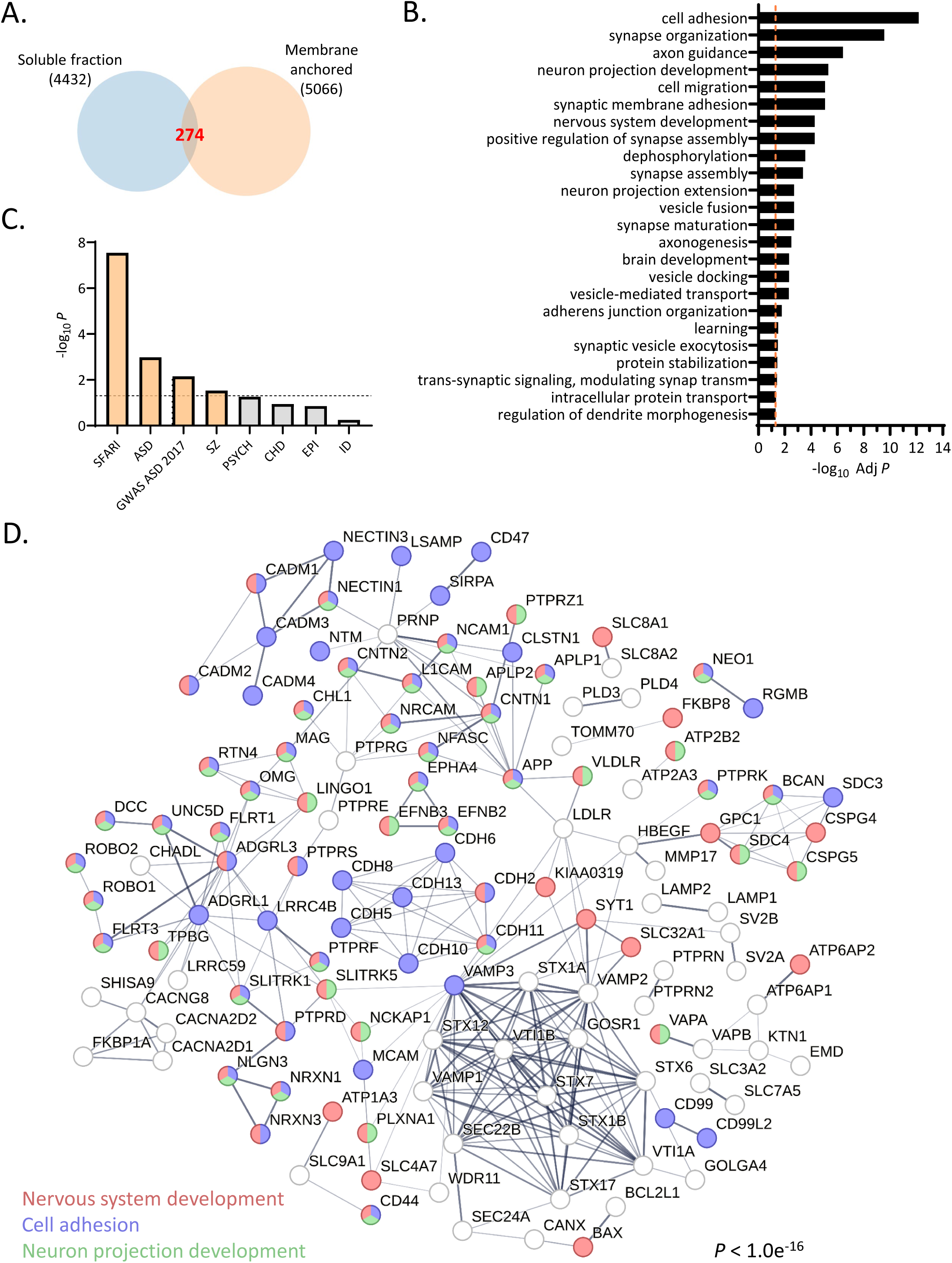
Bioinformatic analysis of the brain sheddome. **A.** Venn diagram showing the overlap of proteins identified in the soluble fraction of control animals (4432 total) with its membrane-anchored subset (274 proteins). Proteins detected in at least 3 of the 6 samples were considered as “present” in the sheddome. n=6, 15-day-old mice (3 females and 3 males). **B.** Gene Ontology (GO) analysis of the 274 proteins in the sheddome. **C.** Enrichment analysis showing disease-relevant gene-encoded proteins within the brain sheddome. **D.** Protein-protein interaction (PPI) network of the brain sheddome, with nodes colored by biological process: red (nervous system development), blue (cell adhesion), and green (neuron projection development).

### Sensory experience differentially modulates the molecular and functional composition of the sheddome in a sex-dependent manner

We next evaluated the impact of whisker trimming (Wtr) on ES by trimming whiskers from birth until 15 days of age every 2-3 days in both male and female mice. This model has been shown to alter neuroplasticity due to sensory deprivation, affecting not only the barrel cortex but also transmodally other cortical regions (45). Analysis of the soluble fraction sheddome, conducted as described previously, revealed that 22 proteins were exclusively detected in the Wtr group, while 10 proteins showed increased levels in this group. We also found 23 proteins that were identified only in the control group, with 7 proteins showing increased levels in the control compared to the Wtr (**Fig.2A and B, Suppl Fig. 1A**). In summary, sensory deprivation induced changes in 62 out of the 392 proteins detected, indicating that 16% of the proteins were affected. GO analysis of the 62 differentially regulated proteins in the sheddome following Wtr demonstrated an enrichment in processes related to phosphorylation, cell migration, monocyte extravasation, and others (**Fig. 2C**).

**Figure 2.**
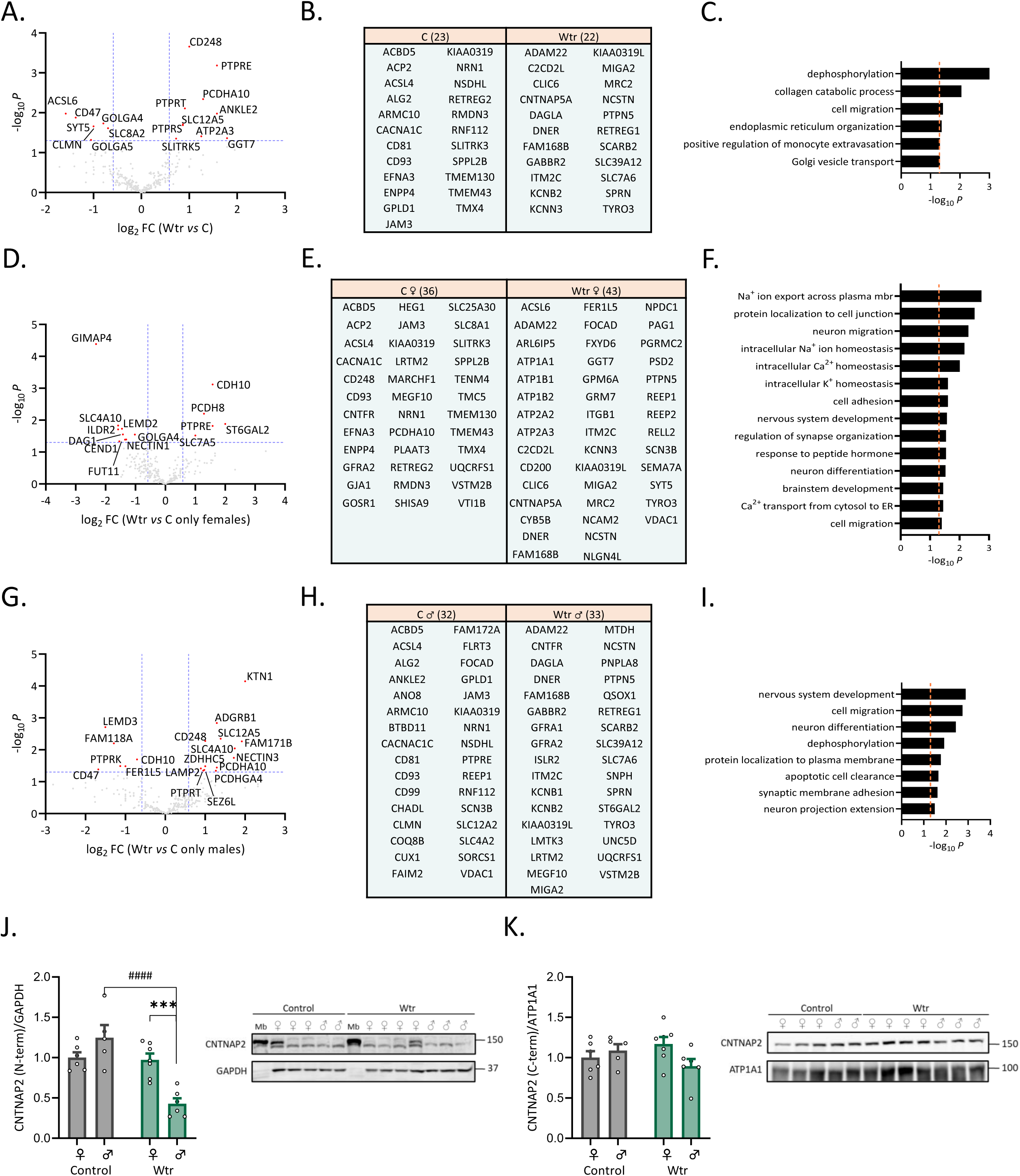
Whisker trimming induces sex-dependent changes in the molecular and functional composition of the sheddome. **A.** Volcano plot of whister trimmed (Wtr) *vs.* control sheddome proteins. **B.** Table of absence/presence. On the left proteins present only in control (23) and on the right proteins present only in Wtr (22). **C.** GO analysis of differentially regulated proteins in the volcano plot plus present/absent proteins. **D.** Volcano plot of Wtr females *vs.* control female mice sheddome proteins. **E.** Table of absence/presence in the female sheddome. On the left proteins present only in control females (36) and on the right proteins present only in Wtr females (43). **F.** GO analysis of differentially regulated proteins in the volcano plot plus present/absent proteins in the female sheddome. **G.** Volcano plot of Wtr males *vs.* control male sheddome proteins. **H.** Table of absence/presence in the male sheddome. On the left proteins present only in control males (32) and on the right proteins present only in Wtr males (33). **I.** GO analysis of differentially regulated proteins in the volcano plot plus present/absent proteins in the male sheddome. In the volcano plots, the dashed lines in the “x” axis correspond to x = -0.585 and x = 0.585 and the dashed line in the “y” axis corresponds to y = 1.3 or *P* = 0.05. Proteins that reached statistical significance are indicated with their names and highlighted in red. Proteins detected in at least 3 of the 6 samples were considered as “present” in the sheddome when n=6 and at least in 2 of three when n=3. **J.** Western blot of mouse soluble fractions from control and Wtr female and male mice, n = 5-7 per group, two-way ANOVA + Holm-Sidak correction, \*\*\**P* < 0.001, ^####^*P* < 0.0001. Data are presented as mean ± SEM. **K.** Western blot of mouse membrane fractions from control and Wtr female and male mice, n=5-7/per group, a two-way ANOVA + Holm-Sidak correction showed no statistical differences between groups. Data are presented as mean ± SEM. Samples were immunoblotted for CNTNAP2 N-term antibody (J) and C-term antibody (K).

We then aimed to analyze whether sensory deprivation impacts the sheddome of female and male mice differentially, given that stronger behavioral changes have been observed in male mice after Wtr in juvenile stages (46). The analysis of the female soluble fraction sheddome showed that 43 proteins were exclusively detected in the whisker trimming group in females, while 5 proteins exhibited increased levels in this group. We also identified 36 proteins that were present in the female control group only, with 9 proteins showing increased levels in this group compared to the female Wtr group. (**Fig. 2D and E**, **Suppl Fig. 1B**). In summary, sensory deprivation induced changes in 93 out of the 392 proteins detected in females, indicating that 24% of the proteins were affected. GO analysis of the 93 differentially regulated proteins in the female mice sheddome following Wtr demonstrated an enrichment in processes such as regulation of intracellular Na^+^, Ca^2+^ and K^+^ levels, nervous system development, neuronal migration, neuron differentiation, among others (**Fig. 2F**).

Finally, the analysis of the male soluble fraction sheddome showed that 33 proteins were exclusively detected in the whisker trimming group in males, while 13 proteins showed increased levels in this group. We also identified 32 proteins that were identified in the male control group only, with 6 proteins showing increased levels in this group compared to the male Wtr group. (**Fig.2G and H, Suppl Fig. 1C**). In summary, sensory deprivation induced changes in 84 out of the 392 proteins detected in the male sheddome, indicating that 21% of the proteins were affected. GO analysis of the 84 differentially regulated proteins in the male mice sheddome after Wtr demonstrated an enrichment in processes shared with the female sheddome, including nervous system development, neuron differentiation, and cell migration, as well as unique processes such as neuron projection extension, synaptic membrane adhesion and protein localization to plasma membrane.(**Fig. 2I**).This data indicates that while the male and female sheddomes share common biological properties, they also differ in specific aspects. We also analyzed the differences between the sheddome in the female Wtr vs the male Wtr, finding 73 differentially regulated proteins, enriched in proteins regulating synaptic membrane adhesion and Na+ homeostasis, among others (**Supp. Fig. 1D-G**). The observation that females exhibit fewer behavioral abnormalities associated with whisker trimming and a greater number of proteins identified in the female sheddome could reflect a more extensive adaptation to the lack of sensory input.

Given the relevance of CNTNAP2 to ASD, particularly its role in regulating dendritic projections and the fact that its levels in the cerebrospinal fluid (CSF) of individuals with ASD are decreased, we analyzed the levels of sCNTNAP2 in the soluble fractions after Wtr. Interestingly, we found that there was a decrease in the levels of sCNTNAP2 only in males, while no changes were observed in the expression levels in the membrane fractions (**Fig. 2J-K**), indicating that the shedding of CNTNAP2 is decreased, independently of the expression levels. The decrease of sCNTNAP2 in males following neuroplasticity changes induced by Wtr, combined with the higher male susceptibility to ASD, reduced sCNTNAP2 levels in the CSF of individuals with ASD, and CNTNAP2’s role in dendritic projection regulation, highlights the potential significance of these findings in elucidating molecular mechanisms that may contribute to altered neuronal connectivity and ASD-associated behavioral phenotypes. These findings suggest that sex-specific regulation of sCNTNAP2 may contribute to the increased ASD prevalence in males, pointing to sCNTNAP2 as a potential target for addressing this sex-based vulnerability.

### Neurodevelopmental regulation of CNTNAP2 ES

To investigate how neuronal maturation affects CNTNAP2 ES, we extracted proteins from extracellular media (ECM) and whole cell lysates (WCL) of cultured cortical neurons at 1, 2, 3 and 4 weeks *in vitro* (WIV). Using both an extracellular (N-terminal) and an intracellular (C-terminal) antibody (**Fig. 3A**), we identified the cleaved ectodomain. Over time, we observed a significant increase in the amount of sCNTNAP2 in the ECM (**Fig. 3B-C**, left panel), which correlated with CNTNAP2 expression levels in the WCL (**Fig. 3C**, middle panel). However, the ratio of sCNTNAP2 to CNTNAP2 expression in the lysates remained consistent across the different weeks in culture (**Fig. 3C**, right panel). These results suggest that as cultured neurons mature and increase dendritic arborization, they release more sCNTNAP2 as a result of higher CNTNAP2 expression levels.

**Figure 3.**
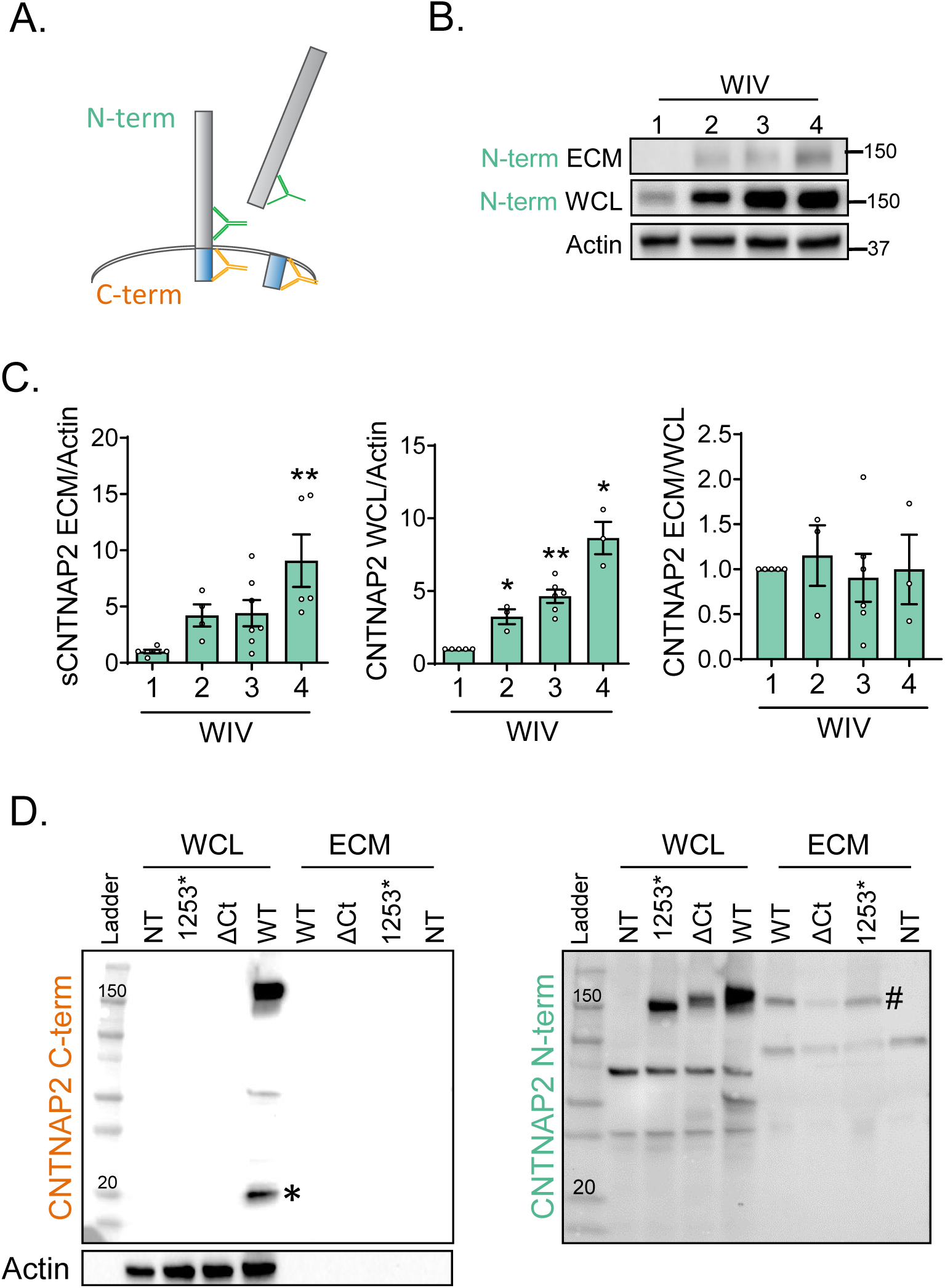
sCNTNAP2 levels increase with time in culture. A. Illustration of antibody recognition sites within CNTNAP2; C-terminal (C-term, orange) antibody recognizes its intracellular domain, while the N-terminal (N-term, green) recognizes its extracellular domain. B. Representative blots of CNTNAP2 in the extracellular media (ECM), whole cell lysates (WCL), and actin in the WCL in dissociated rat neurons after 1, 2, 3, 4 weeks *in vitro* (WIV). C. Analysis of CNTNAP2 levels in the ECM and WCL with time *in vitro*. CNTNAP2 levels increase with time in culture in the ECM (left panel) and the WCL (middle panel). No differences are observed in the ratio of CNTNAP2 in ECM *vs* WCL (right panel). Data are presented as mean ± SEM. <u>ECM data</u>: n = 3-7, Kruskal-Wallis test + Dunn’s post hoc, **P=0.0061; <u>WCL data:</u> n = 3-6, Mann-Whitney test, 1 *vs* 2 WIV * *P* =0.0490, 1 *vs* 3 WIV \*\**P* = 0.012, 1 *vs* 4 WIV \**P* = 0.0168. D. Whole cell lysates (WCL) and extracellular media (ECM) from HEK293 cultures transfected with WT, ΔC-term-CNTNAP2 (ΔCt) and the Cortical Dysplasia-Focal Epilepsy Syndrome (CDFE)-associated variant 1253*. NT, non-transfected. Samples were immunoblotted for CNTNAP2 C-term antibody (left) and N-term antibody (right). *Indicates the intracellular C-terminal fragment produced after ectodomain shedding. # indicates the released CNTNAP2 ectodomain.

A rare variant CNTNAP2 1253*, has been reported in Old Order Amish family members with a syndromic form of autism called Cortical Dysplasia Focal Epilepsy (CDFE) syndrome (47). Individuals with this variant exhibit neurodevelopmental abnormalities, including language regression, hyperactivity, impulsive and aggressive behavior, autism, and intellectual disability. To evaluate whether the 1253* variant produces a soluble ectodomain, we transfected HEK293 cells with CNTNAP2 WT, the 1253* variant, and a truncated version of CNTNAP2 lacking the intracellular domain (ΔCt). Interestingly, all constructs showed a band at around 150 kDa when using the extracellular N-terminal antibody (**Fig. 3D**, right panel, marked with a pound symbol), which was absent in the not transfected cells and not detected when using the intracellular C-terminal antibody (**Fig. 3D**, left panel). These findings demonstrate that the disease-associated variant 1253* releases a soluble form of CNTNAP2 that cannot regulated by ES. Instead, it would be released through the secretory pathway, lacking the standard ES regulatory control. Additionally, the data show that the C-terminal domain is not required for CNTNAP2 shedding, as the ΔCt construct still released sCNTNAP2 into the ECM.

Interestingly, the blots in Fig. 3D also showed the presence of a band at approximately 20 kDa in the WCL when using the intracellular antibody (**Fig. 3D**, marked with a star), a band that was missing in the CNTNAP2 ΔCt and the 1253* constructs. The 1253* variant lacks the intracellular and transmembrane domains, as well as part of the extracellular domain, while the last 16 amino acids on the N-terminal side differ from the WT version of CNTNAP2. The 20 kDa band was also absent in the ECM samples, in non-transfected cells (**Fig. 3D**, left panel), and when using the extracellular antibody (**Fig. 3D**, right panel). These findings suggest that CNTNAP2 ES generates an intracellular fragment of about 20 kDa. In summary, these results demonstrate that neuronal maturation is associated with increased levels of sCNTNAP2 due to increased CNTNAP2 expression. Furthermore, the disease-associated variant 1253* releases a soluble form of CNTNAP2, through the ES-independent secretory pathway bypassing the usual regulatory mechanisms of ES. This highlights how variations in CNTNAP2 can disrupt its regulation and release, with potential impacts on its neurodevelopmental functions.

### sCNTNAP2 increases dendritic arborization

After observing that sCNTNAP2 levels increase as neurons mature and increase their dendritic complexity in culture, we investigated its contribution to dendritic arborization. We transfected three-week-old rat cortical neurons with GFP and then treated them with purified sCNTNAP2 at 10 nM for 24h. We then analyzed dendritic arborization by Sholl analysis. Sholl analysis revealed that sCNTNAP2 significantly enhanced dendritic branching in both excitatory and inhibitory neurons compared to the vehicle control (**Fig. 4A-B).**

**Fig. 4.**
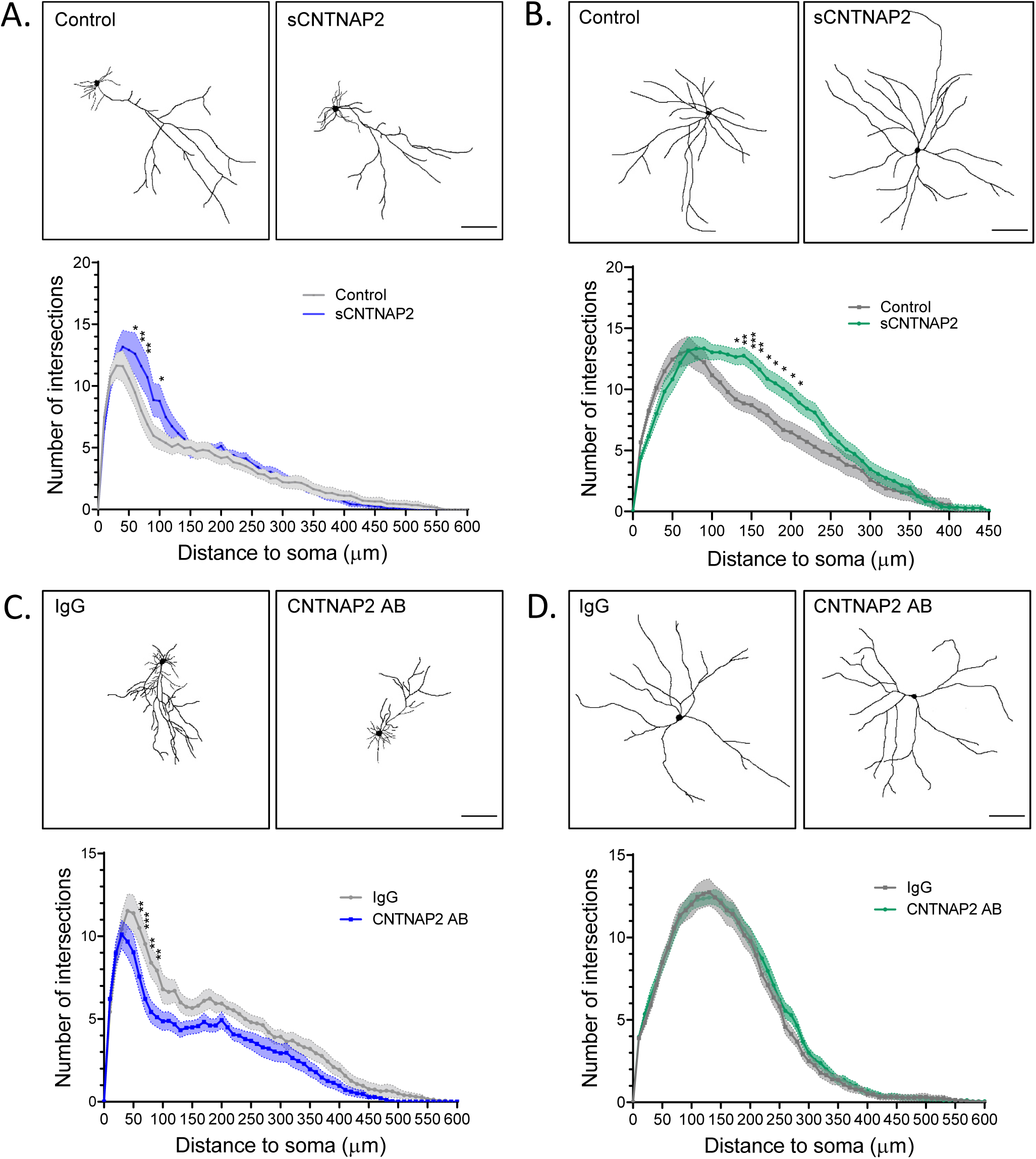
sCNTNAP2 incubation increases dendritic arborization. Top panel shows traces of dendritic arbors of GFP-transfected excitatory neurons treated with 10 nM sCNTNAP2 or vehicle as a control. Dendritic trees were measured via Sholl analysis. N = 3 cultures, n = 23-28 neurons per group. A two-way ANOVA analysis showed significant effect of genotype (*P* = 0.0009) and genotype–distance interaction (*P* = 0.0097). Individual data points that were significant after Holm-Sidak correction are indicated: \**P* < 0.05; \*\**P* < 0.01. **B.** Traces of the dendritic arbors of GFP-transfected inhibitory neurons treated with 10 nM sCNTNAP2 or vehicle. N = 3 cultures, n = 27-28 neurons per group. Dendritic trees were measured via Sholl analysis. N = 3 cultures, n = 23-28 neurons. A two-way ANOVA analysis showed significant effect of genotype (*P* < 0.0001) and genotype–distance interaction (*P* < 0.0001). Individual data points that were significant after Holm-Sidak correction are indicated: \**P* < 0.05; \*\**P* < 0.01, \*\*\**P* < 0.001. All data are presented as mean ± SEM. Scale bar, 100 μm. **C.** Top panel shows traces of dendritic arbors of GFP-transfected excitatory neurons treated with a control IgG antibody or the N-terminal CNTNAP2 antibody (CNTNAP2 AB). Dendritic trees were measured via Sholl analysis. N = 3 cultures, n = 28-30 neurons. A two-way ANOVA analysis showed significant effect of genotype (*P* < 0.0001) and no genotype–distance interaction (*P* = 0.1005). Individual data points that were significant after Holm-Sidak correction are indicated: \*\**P* < 0.01, \*\*\**P* < 0.001. **D.** Top panel shows traces of dendritic arbors of GFP-transfected inhibitory neurons treated with a control IgG antibody or the CNTNAP2 AB. Dendritic trees were measured via Sholl analysis. N = 3 cultures, n = 29-31 neurons. A two-way ANOVA + Holm-Sidak correction showed no statistical differences.

To further explore the role of the CNTNAP2 ectodomain in dendritic architecture, we treated neuronal cultures with either an N-terminal anti-CNTNAP2 antibody or an IgG control for 24 hours. Consistent with our previous findings, we observed a significant reduction in dendritic branching of pyramidal neurons (**Fig. 4C**), although there was no effect on inhibitory neurons (**Fig. 4D**). These results suggest that sCNTNAP2 plays a crucial role in promoting dendritic branching, highlighting its potential as a key regulator of neuronal connectivity and plasticity and its potential significance in neurodevelopment.

### CNTNAP2 mutations shed different levels of CNTNAP2 ectodomain

CNTNAP2 undergoes MMP9-dependent ES, leading to the release of sCNTNAP2 into the CSF (7). Notably, ASD patients show decreased sCNTNAP2 levels in this fluid (7) and as we have described sCNTNAP2 increases dendritic branching, suggesting that alterations in sCNTNAP2 may impact neuronal function and contribute to disease.

To examine how CNTNAP2 variants affect sCNTNAP2 levels, we focused on ASD-associated mutations N418D, R1119H, D1129H, T1278I, and 1253* (29) (**Fig. 5A**). Following a 2-day transfection of HEK293T cells with CNTNAP2 WT and these variations, we found that R1119H, D1129H, and T1278I exhibited reduced ES into the ECM (**Fig. 5B**). Interestingly the 1253* variant showed similar sCNTNAP2 levels compared to WT.

**Figure 5.**
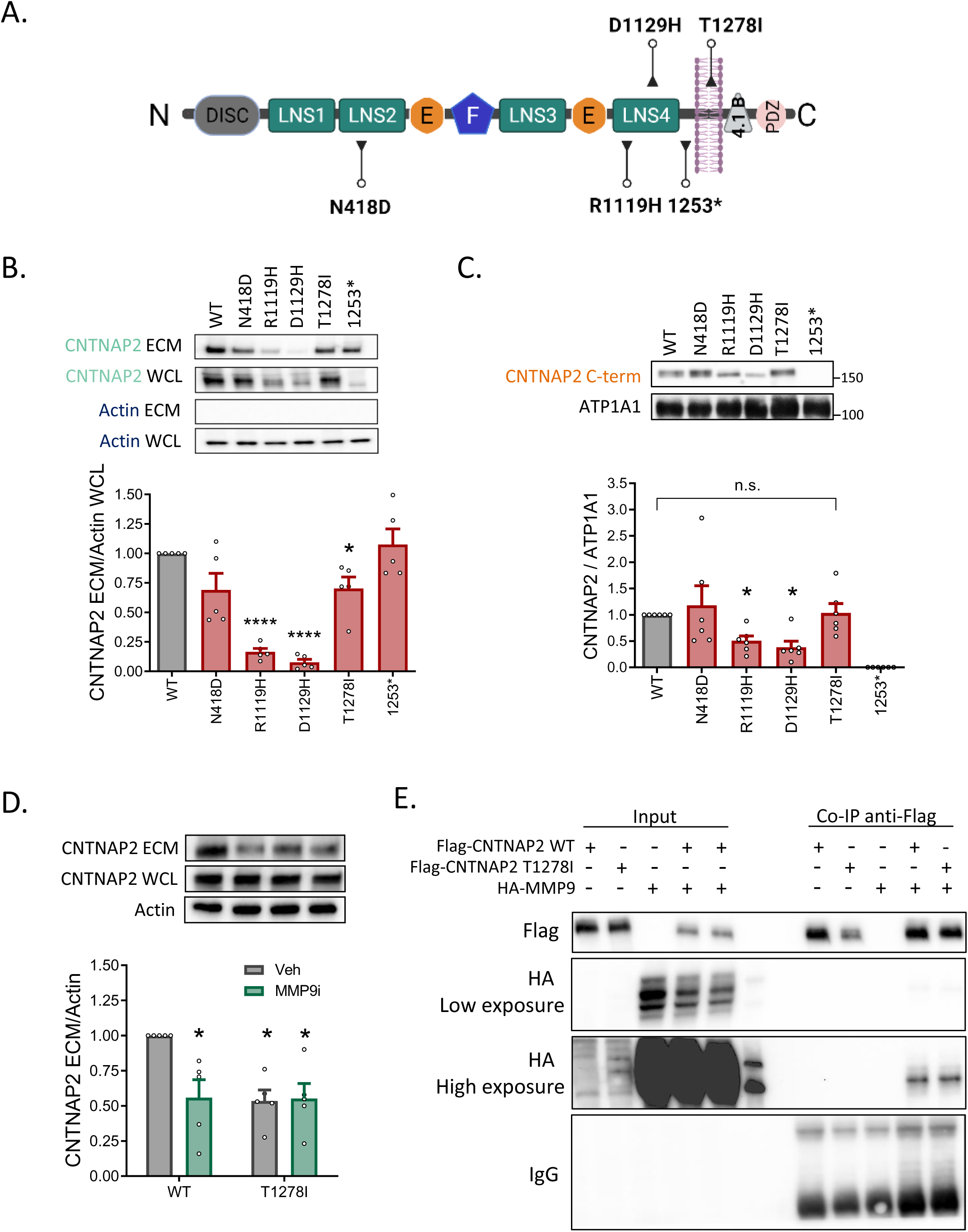
ASD-related genetic variation on CNTNAP2 leads to decreased levels of shed ectodomain. **A.** CNTNAP2 domain structure and location of ASD-related mutations assessed in this study (N418D, R1119H, D1129H, T1278I and 1253*). CNTNAP2 possesses the following domains: “DISC”, discoidin domain; four “LNS”: laminin G domains, two “E”: epidermal growth factor-like domain; “F”: fibrinogen-like domain; “4.1B”, 4.1 binding domain; “PDZ”, PDZ binding domain. **B.** Representative blot and quantification of CNTNAP2 released into the extracellular media (ECM) measured by Western blot (WB) in HEK293T cells transfected with CNTNAP2 WT and 5 ASD-associated variations. n = 5, paired *t*-test compared to WT group. \**P* = 0.0037, \*\*\*\**P* < 0.0001. **C.** Representative blot and quantification of CNTNAP2 levels in membrane fractions obtained from HEK293T cells transfected with CNTNAP2 WT and 5 ASD-associated variations. n = 6, paired *t*-test compared to WT group. \**P* < 0.05; n.s., not significant. **D.** Representative blot and WB analysis of sCNTNAP2 in the ECM in WT and T1278I transfected HEK293T cells. (n = 5; 2-way ANOVA + Šídák’s multiple comparisons test: WT *vs.* WT+MMP9 inh, *P* = 0.0212; WT *vs.* T1278I, *P* = 0.0143; WT *vs.* T1278I+MMP9 inh, *P* = 0.0181. All data are shown as mean ± SEM. **E.** Coimmunoprecipitation experiment of HEK293 cells transfected or not with CNTNAP2 WT, CNTNAP2 T1278I and MMP9.

Previous studies indicate that R1119H and D1129H are misfolded and retained in the endoplasmic reticulum, leading to decreased protein expression (48, 49). To assess whether these CNTNAP2 variants reached the membrane for shedding, we extracted membranes from cell lysates. Western blot analysis confirmed that R1119H and D1129H had decreased levels at the membrane, while N418D and T1278I showed no significant differences compared to the WT (**Fig. 5C**). Therefore, the reduced shedding of CNTNAP2 in the T1278I variant is not due to decreased membrane expression.

To investigate the underlying cause of decreased sCNTNAP2 levels in the T1278I variant, we incubated CNTNAP2 WT and T1278I-transfected HEK293T cells with MMP9 inhibitor I (MMP9i) at 1 μM overnight. While the inhibitor reduced ES by nearly 50% in WT cells, it had no effect on T1278I-transfected cells (**Fig. 5D**). This suggests that the lack of ES in CNTNAP2 T1278I may result from a failure to be regulated by MMP9.

To further understand this, we conducted co-immunoprecipitation experiments after overexpressing MMP9 and either WT or T1278I CNTNAP2 in HEK293T. We found that co-immunoprecipitated levels of MMP9 were comparable between T1278I mutants and WT (**Fig. 5E**), indicating that while T1278I can physically interact with MMP9, it is resistant to shedding. Further experiments are needed to clarify why MMP9 fails to cleave CNTNAP2 T1278I.

### Disease-relevant mutations on CNTNAP2 impact dendritic complexity

Previous studies have established that CNTNAP2 plays a critical role in regulating dendritic architecture (33, 34). However, the impact of ASD-related CNTNAP2 mutations on dendritic arborization and soma size remains unclear. To investigate the effects of these ASD-associated variations on neuronal morphology, we transfected three-week-old rat cortical neurons with CNTNAP2 WT, T1278I, and 1253*, along with GFP for cell filling, and maintained the cultures for three days. Following immunostaining for GFP, we performed Sholl analysis to quantify dendritic intersections at 10-μm increments. Notably, the T1278I variant resulted in significantly fewer dendritic intersections in cortical pyramidal neurons compared to WT neurons (**Fig. 6A**), while showing no significant impact on inhibitory neurons (**Fig. 6B**). In contrast, neurons expressing the 1253* mutation displayed enhanced dendritic branching in both pyramidal and inhibitory neuron populations (**Fig. 6A-B**). We assessed the morphology of neuron somas following transfection with CNTNAP2 WT and the variants; no significant changes in the size of either pyramidal or inhibitory neuron somas were observed (**Fig. 6C-D**). Collectively, these results demonstrate that ASD-related mutations in CNTNAP2 distinctly influence dendritic branching in cultured cortical neurons, suggesting a potential mechanism through which these variations may contribute to the neurodevelopmental differences observed in ASD.

**Figure 6.**
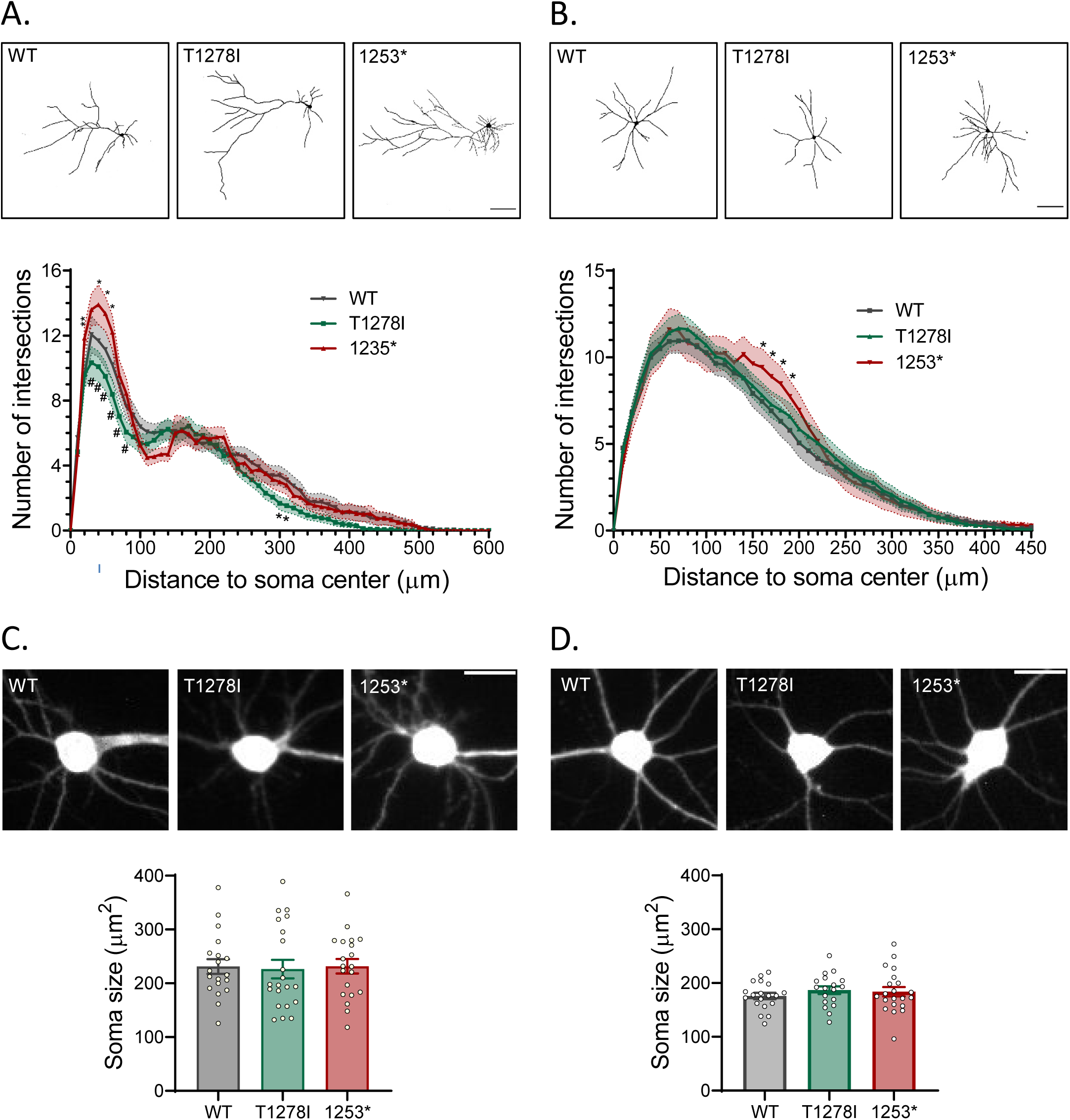
ASD-associated mutations on CNTNAP2 impact dendritic complexity. **A.** Top panel: Representative traces of CNTNAP2 WT, T1278I and 1253* pyramidal neurons used in analyses. Lower panel: Sholl analysis of total dendrites in CNTNAP2 WT (N = 4 cultures, n = 28 neurons), T1278I (N = 4 cultures, n = 34 neurons) and 1253* (N = 2 cultures, n= 20 neurons). A two-way ANOVA showed a significant effect of genotype (*P* < 0.0001) and genotype– distance interaction (*P* = 0.0174). Individual data points that were significant after Holm-Sidak correction are indicated: \**P* < 0.05; \*\**P* < 0.01. Scale bar 100 μm. **B.** Top panel: Representative traces of CNTNAP2 WT, T1278I and 1253* transfected inhibitory neurons used in analyses. Lower panel: Sholl analysis of total dendrites in CNTNAP2 WT (N = 4 cultures, n = 33 neurons), T1278I (N = 4 cultures, n = 37 neurons) and 1253* (N = 2 cultures, n = 19 neurons); significant effect of genotype (*P* = 0.0450) and no genotype–distance interaction (*P* > 0.9999). Individual data points that were significant after Holm-Sidak correction are indicated: \**P* < 0.05. Scale bar 100 μm. **C.** Representative images and analysis of soma sizes of CNTNAP2 WT, T1278I and 1253* transfected excitatory neurons (N=2 cultures, n=19-21 neurons). Scale bar 20 μm. **D.** Representative images and analysis of soma sizes of CNTNAP2 WT, T1278I and 1253* transfected inhibitory neurons (N=2 cultures, n=18-21 neurons). Scale bar 20 μm. All data are presented as mean ± SEM.

## DISCUSSION

### Sex-Specific Alterations in Ectodomain Shedding and Their Consequences for Neuronal Connectivity in ASD

In recent years, the role of ES in brain disorders is gaining increasing attention. It has been hypothesized that ES could be implicated in neurodevelopmental conditions, including ASD (7), though more evidence is required to substantiate this hypothesis. ASD is characterized by distinct alterations in synaptic plasticity, brain connectivity, and neural circuitry, with a notable prevalence in males compared to females. This highlights the need to study how sex impacts the molecular mechanisms underlying ASD. This is highly relevant because identifying therapeutic strategies that specifically target the neurobiological pathways affected in males and females could potentially lead to more effective treatments. Interestingly, in this study we have analyzed the molecular and functional composition of the brain sheddome at 15 days of age in mice, identifying 274 proteins that could undergo ES. Interestingly, the proteins in the cortical sheddome are enriched in key processes for neurodevelopment such as neuronal cell migration, cell adhesion, axonogenesis, axon guidance, neuron projection development, dendrite morphogenesis, synapse assembly and maturation among others. Not surprisingly, the cortical sheddome is enriched in risk factors for ASD, supporting previous data (7). Moreover, ASD-associated CNTNAP2 variants impair its regulation by ES, impacting dendritic arborization —a key feature for maintaining adequate brain connectivity and synaptic plasticity, both crucial processes that are disrupted in ASD. This dysregulation can contribute to the connectivity deficits commonly observed in ASD, as inadequate dendritic complexity may hinder proper synaptic formation and neural network integration.

We observed that sensory deprivation led to a decrease in sCNTNAP2 levels only in male cortices, while previous studies have reported reduced sCNTNAP2 levels in the CSF of individuals with ASD (7). This finding is particularly significant, as sCNTNAP2 has been shown to regulate neuronal synchrony (7) and, in our study, to enhance dendritic branching—processes that are essential for proper circuit formation and function. Collectively, these sex-specific reductions in sCNTNAP2 may exacerbate the developmental and cognitive challenges frequently associated with ASD, particularly in males.

Furthermore, our findings suggest that sCNTNAP2 promotes dendritic branching, thereby contributing to CNTNAP2’s regulation of neuronal plasticity and connectivity. This role of ES in modulating connectivity aligns with other studies demonstrating that the ectodomains of various cell adhesion molecules—such as NCAM (Neural Cell Adhesion Molecule), L1CAM (L1 Cell Adhesion Molecule), neurexin-1β, CHL1 (Close Homolog of L1), NEGR1 (Neurite Growth Promotion Associated Protein 1), and NRG1 (Neuregulin 1) (50–54)—positively modulate neurite branching. Future research should address if shed ectodomains impact dendritic arborization in a sex-specific manner.

### Mechanisms of disrupted ES in ASD-associated CNTNAP2 variants

We found that the levels of sCNTNAP2 are altered in ASD-associated variations of CNTNAP2. The variants R1119H and D1129H exhibited decreased levels of the shed ectodomain, likely due to the mutations causing misfolding of the protein, which results in retention in the endoplasmic reticulum and subsequent degradation, as previously described (48). This is similar to a decrease in expression and proper localization of the protein at the plasma membrane, where ectodomain shedding (ES) of CNTNAP2 occurs. In contrast, we and others have shown that the T1278I mutant appropriately reaches the plasma membrane (48), and bound MMP9; however, it was not cleaved by it, as treatment of T1278I-transfected HEK cells with an MMP9 inhibitor did not modify sCNTNAP2 levels in the extracellular media. This mutation, although located within the transmembrane domain, likely induces conformational changes that permit the CNTNAP2-MMP9 interaction, while blocking shedding. Finally, the CNTNAP2 1253* mutant, associated with CDFE (47), displayed similar levels of sCNTNAP2 as the WT. The mechanism regulating its release into the extracellular media differs for this mutant, as it is translated as a soluble protein from the beginning. Consequently, this mutant is constitutively and continuously secreted, unlike the activity-regulated ES. Remarkably, this modification in the processing of sCNTNAP2 may have implications for connectivity, as we found that dendritic arborization is increased in neurons transfected with the 1253* mutant. In summary, we observed three mechanisms associated with decreased shedding of CNTNAP2, including misfolding and retention in the endoplasmic reticulum of variants R1119H and D1129H, conformational changes in the T1278I variant that block shedding despite proper binding to MMP9, and the continuous secretion of the 1253* mutant as a soluble protein, which differs from activity-regulated shedding.

### Impact of CNTNAP2, CNTNAP2 ASD-associated variations and sCNTNAP2 on connectivity in ASD

The finding that sCNTNAP2 enhances dendritic branching adds to the growing body of evidence regarding CNTNAP2’s role in regulating neuronal plasticity and connectivity. Studies on *Cntnap2* knockout (KO) mice have demonstrated reduction in dendritic branching in inhibitory but not in excitatory neurons, both *in vitro* and *in vivo* (34). Similarly, deficits in arborization have also been reported in cortical excitatory neurons after *Cntnap2* knock down (33). In our study, however, we found a more consistent phenotype on excitatory neurons: We found a modification in excitatory neuron branching when treating with sCNTNAP2 and when blocking the extracellular domain of CNTNAP2 with an N-terminal antibody. Conversely, in inhibitory neurons, we only observed an effect when treating the cells with sCNTNPA2, but not with the blocking antibody. Moreover, CNTNAP2 T1278I and CNTNAP2 1253* modulated excitatory neuron arborization, but in inhibitory neurons only the 1253* variant induced changes. This apparent discrepancy in findings may be attributed to the duration of CNTNAP2 modification, which is chronic in the KO model, but acute in treatments with sCNTNAP2, the blocking antibody, KD, or in the mutants. This suggests that while inhibitory neurons may exhibit a degree of resilience to the loss of CNTNAP2 function, the long-term consequences can be detrimental. This model may better represent the condition of patients with CNTNAP2 loss of function. Furthermore, our data indicates that although excitatory neurons could be initially more sensitive to CNTNAP2 loss, they may adapt over time.

Apart from its role on dendritic arborization, previous studies have shown that CNTNAP2 positively regulates axonal outgrowth in neurons at DIV3. Additionally, CNTNAP2 variants R1119H and N407S exhibit a dominant-negative effect on axonal growth, whereas I869T and G731S variants do not rescue axonal growth defects in heterozygous *Cntnap2* KO neurons (49). Collectively, these findings underscore the essential role of CNTNAP2 in structural connectivity among cortical neurons, potentially contributing to the pathogenesis of ASD.

### Potential Therapeutic Pathways

Understanding the proteolytic regulation of CNTNAP2 and its impact on connectivity could lead to targeted interventions. For instance, modulating MMP9 or other enzymes involved in shedding or treatment with CNTNAP2 could represent a therapeutic target for ASD, particularly for variants with defective CNTNAP2 ES.

## MATERIALS AND METHODS

### Animals

All animals were kept in a controlled environment with a photoperiod of 08:00– 20:00 (light) and a temperature maintained at 22 ± 1°C, with unlimited access to standard food and water. The experiments were conducted in accordance with either local and European regulations (directive 2010/63/EU) and received approval from the Ethical Committee of Universidad Complutense de Madrid (ref. PROEX 305.6/22); or with the approval of the Institutional Animal Care and Use Committee (IACUC) at Northwestern University.

### Antibodies, plasmids, and chemicals

For Western blot analysis the following primary antibodies were used: CNTNAP2 (N-terminal, Neuromab, cat# 75–075), CNTNAP2 (C-terminal, Millipore, cat# AB5886), Flag (SigmaAldrich, cat# F1804), and anti-HA (Abcam, cat# ab20084). Secondary antibodies included: goat polyclonal anti-mouse IgG (SigmaAldrich, cat# A4416) and goat polyclonal anti-rabbit IgG (ThermoFisher, cat# 31460). For detecting transfected cells and conducting Sholl analysis we used the anti-GFP antibody (Abcam, cat # ab13970).

The pEGFP-N2 plasmid was purchased from Clontech (Mountain View, CA, USA). CNTNAP2 plasmids used for HEK293T cell and neuronal transfections, including WT, N418D, R1119H, D1129H, T1278I, and 1253* were kindly donated by Prof. Davide Comoletti (48).

The Flag-CNTNAP2 plasmid was generated by subcloning human CNTNAP2 cDNA (gift from Dr. Elior Peles, Weizmann Institute of Science, Israel) into the pEGFP-N2 vector (restriction sites BamHI and NotI, removing EGFP) as previously described (55). MMP9 inhibition experiments used MMP9 inhibitor I (Calbiochem) at a concentration of 1 μM.

For the experiments designed to block N-terminal CNTNAP2, two antibodies were incubated for 24h at 2 μg/well (800 μL media per well in 12 welll plates): mouse CNTNAP2 N-terminal (Neuromab, cat# 75–075) and. mouse IgG control antibody (Abcam, cat# ab376355).

### Sensory deprivation by whisker-trimming

This procedure was performed according to a protocol published elsewhere (45), with slight modifications. Briefly, CD1 male and female mice were randomly assigned to either the control group or the whisker trimming (sensory deprivation) group. All pups were anesthetized with isoflurane to achieve the appropriate level of sedation before each trimming session, ensuring consistent handling across groups. In the sensory deprivation group, all whiskers were carefully trimmed, while in the control group, pups were similarly handled but without any whisker trimming. The trimming procedure was repeated every 2–3 days from postnatal day 0 (P0) to postnatal day 15 (P15), after which the mice were euthanized for experimental analysis.

### Isolation of membrane and soluble fractions from mouse cortex and bioinformatic identification of the cortical sheddome

On postnatal day 15, mice were sacrificed by decapitation, and their cerebral cortices were dissected. The cortices were mechanically homogenized using a glass homogenizer in a tris-based buffer (TS) containing 50 mM tris-HCl, 150 mM NaCl, and a protease inhibitor cocktail (ThermoFisher), with 6.8 mL of buffer per gram of cortical tissue. The homogenate was centrifuged at 1,500 g for 10 min at 4°C to remove nuclei, cell debris, and other large organelles. The resulting pellet was then discarded, and the supernatant was ultracentrifuged at 100,000 g for 2 hours at 4°C with a 100 Ti rotor (Beckman Coulter) in a Beckman XL-90 ultracentrifuge, separating the membrane (P2) and soluble (S2) fractions. The P2 fraction was resuspended in TS buffer containing 1% triton X-100, 0.5% sodium deoxycholate, and 1% sodium dodecyl sulfate (SDS). Both P2 and S2 fractions were subsequently analyzed by Western blotting. Additionally, the S2 fraction was analized by liquid chromatography tandem mass spectrometry (LC-MS/MS) for molecular and functional characterization of the brain’s soluble fraction sheddome, following the methodology previously described (8). To identify the subset of proteins undergoing ES identified in the soluble fractions, we bioinformatically selected proteins that contain at least one transmembrane domain or glycosylphosphatidylinositol (GPI) anchor based on UniProt annotations, as previously described (8).

### Liquid chromatography with tandem mass spectrometry

Samples prepared for analysis by liquid chromatography coupled with tandem mass spectrometry (LC-MS/MS) were processed using the PreOmics iST kit, following the manufacturer’s instructions. Briefly, samples were diluted in cold acetone, centrifuged, and the supernatant was discarded. Each precipitated sample was treated with Lyse buffer at 95°C for 10 min, transferred to a column, and Digest solution was added, followed by incubation at 37°C for 2 hours. Stop buffer was then added, and samples were centrifuged at 3,800 g for 2 min. The resulting digest was washed with Wash 1 and Wash 2 buffers and eluted twice with Elute buffer. Samples were dried in a Speed-Vac (Thermo-Savant) and reconstituted in LC-load. Peptide concentration was determined using the Qubit system (ThermoFisher), calculating the amount needed to inject 0.5 μg into the nanoHPLC (ThermoFisher).

From the digested peptide mixture, 0.5 μg was injected into the Easy-nLC 1000 nano-HPLC, concentrated on a PEPMAP100 C18 NanoViper Trap precolumn (Thermo Fisher), and separated on a 50 cm PEPMAP RSLC C18 column (Thermo Fisher) using a 2% to 40% ACN and 0.1% formic acid gradient over 120 minutes.

Chromatographically separated peptides were electrospray ionized in positive mode and analyzed on a Q Exactive HF mass spectrometer (ThermoFisher) in data-dependent acquisition (DDA) mode. MS scans were performed between 350 and 1,700 Da, selecting the 10 most intense precursors (with charges between 2+ and 6+) for high collision energy dissociation (HCD) and acquiring the corresponding MS/MS spectra.

Data from the shotgun analysis were processed using the Proteome Discoverer software (ThermoFisher). Peptide spectrum matches (PSMs) of each MS/MS spectrum were identified by comparison with theoretical mass lists from the mouse protein database in the UniProt repository using the Sequest search engine. Peptides were assigned to their corresponding proteins by applying the principle of parsimony to generate a “Master” protein when a peptide could be associated with multiple proteins. The Percolator algorithm was used to estimate the false-positive rate (FDR), filtering identifications with high confidence with a q-value <0.01.

### Biological process enrichment analysis

Gene annotation enrichment analysis was performed using the DAVID v2023q4 tool (56), focusing on gene ontology (GO) terms related to biological processes. To ensure consistency in nomenclature, all identifiers were translated to entrez gene IDs. GO analysis of the soluble fraction sheddome was conducted using all membrane-anchored proteins (both transmembrane and GPI-anchored) annotated by UniProt as a background to eliminate any potential bias toward this group of proteins. The -log_10_ of the *P*-values were calculated and plotted for non-redundant categories of GOTERM_BP_DIRECT, with a significance threshold set at a value of 1.3.

### Disease enrichment analysis

A hypergeometric test was used to calculate the probability of finding an over-enrichment of disease-relevant genes in the secretome and CSF gene ID lists.

For disease-relevant gene lists, we included *de novo* gene lists (ASD: autism spectrum disorder, SZ: schizophrenia, PSYCH: psychiatric disorders, CHD: congenital heart disease, EPI: epilepsy, ID: intellectual disability) from the supplemental data in (43), since *de novo* exonic mutations link specific genes to particular disorders. Only *de novo* variants affecting protein-coding regions or splice sites were included (e.g., missense, frameshift, splice site mutations, altered stop or start codons, insertions, deletions). Additionally, ASD risk factors from the SFARI database and from a GWAS meta-analysis of individuals with autism (44) were incorporated.

### Protein-protein interaction network analysis

Protein–protein interaction (PPI) network analysis was performed using STRING v12.0. The network edges indicate the confidence of PPIs, based on experimental data and databases as the primary sources of interaction, with a medium confidence level. The k-means clustering method was applied to refine the network.

### Western blotting

Western blotting was performed to analyze protein expression levels. The protein concentration of each sample was determined using the bicinchoninic acid method (Pierce BCA Protein Assay Kit, ThermoFisher). 2X sample buffer (BioRad, cat#1610737EDU) and β-mercaptoethanol (Sigma-Aldrich) were then added to each sample, followed by a 2 min boiling step at 95°C in a thermoblock. Electrophoresis was carried out at 120V, after which the proteins were transferred onto polyvinylidene fluoride (PVDF) membranes pre-activated in methanol. The membranes were then blocked with 3% bovine serum albumin (BSA) prepared in TBS-tween buffer (19 mM Tris, 137 mM NaCl, 2.7 mM KCl, and 0.1% tween20). Finally, the membranes were incubated with specific primary and secondary antibodies and visualized using SuperSignal West Pico PLUS (BioRad), a chemiluminescent substrate for horseradish peroxidase. Luminescence signals were captured using a ChemiDoc apparatus and analyzed with the ImageLab sofware (BioRad).

### HEK293T culture and transfection

HEK293T cells (ATCC) were cultured at 37°C and 5% CO_2_ in Dulbecco’s modified Eagle’s medium (DMEM, ThermoFisher) containing 10% fetal bovine serum (Corning) and penicillin/streptomycin (SigmaAldrich). They were passed twice per week during the linear growth phase to maintain sub-confluent cultures. Cells were transfected using Lipofectamine 2000 (ThemoFisher, cat# 11668019) following the manufacturer’s instructions. Experiments were performed 48 h post-transfection.

### Precipitation of extracellular media proteins by TCA method

Supernatants were collected, centrifuged for 3 min at 16,000 rpm to eliminate any potential cell debris and then 0.015% deoxycholate was added to the supernatants. After a 5-min incubation, trichloroacetic acid (TCA) was added to the supernatants to achieve a final concentration of 20%. After 1 h on ice, the samples were centrifuged at 18,000 g. The pellets were washed three times with ice-cold acetone. The acetone was evaporated at room temperature, and the pellets were dissolved in 2X sample buffer (BioRad, cat# 1610737EDU).

### Extraction of membrane fractions from HEK cells

Extraction of membrane fractions from HEK293T cells was performed as previously described (57). Briefly, transfected HEK293T cells were homogenized in cold sucrose buffer (20 mM HEPES pH 7.4, 320 mM sucrose, 5 mM EDTA) supplemented with protease inhibitor cocktail (ThermoFisher). Homogenates were centrifuged at 3,000 g for 10 min at 4°C to pellet nuclei, large organelles, and cell debris. The resulting supernatants (S1) were then centrifuged at 21,000 g for 1 h at 4°C to obtain a crude membrane pellet (P2), which was resuspended with Tris buffer (50 mM Tris pH 7.4, 150 mM NaCl, 5 mM EDTA, 1% Triton X-100, 0.5% deoxycholate, 0.1% SDS with protease inhibitors) and stored at -20°C until analysis by Western blot was performed.

### Neuronal culture

High density (100,000 cells/cm^2^) cortical neuron cultures were prepared from Sprague-Dawley rat E18 embryos as described previously (58). Briefly, neurons were plated onto coverslips coated with poly-D-lysine (0.2 mg/mL; SigmaAldrich), in feeding medium (Neurobasal medium supplemented with B27 (Invitrogen), 0.5 mM glutamine, and penicillin/streptomycin). Four days later, 200 μM DL-2-Amino-5-phosphonovaleric acid (APV) (Abcam, cat# ab120004) was added to the medium. Half of the feeding medium containing APV was replaced twice a week until the neurons were used.

### Neuronal transfections

Cortical neurons were transfected on day *in vitro* (DIV) 21 using Lipofectamine 2000, following the manufacturer’s recommendations. The neurons were maintained in feeding medium for 3 days post-transfection.

### Immunocytochemistry

DIV 21 neurons were first washed in PBS and then fixed in 4% formaldehyde- 4% sucrose in PBS for 10 min. Then, they were washed 3 times with PBS for 10 min. Fixed neurons were then simultaneously permeabilized and blocked for 1h at room temperature in permeabilization/blocking buffer (0.1% BSA, 4% NGS, 0.1% Triton-X-100 in PBS). Following this, the primary antibody against GFP was applied and incubated overnight at 4°C. The coverslips were washed 3 times with PBS and incubated for 45 min at room temperature with the fluorophore-secondary antibody Alexa Fluor^®^488 (LifeTechnologies). The coverslips were then mounted onto slides using ProLong Antifade reagent (Invitrogen) and stored at 4°C until the acquisition of the images. All images were acquired in the linear range of fluorescence intensity.

### Dendrite analysis

To examine dendritic morphology, neurons expressing GFP were imaged using a Zeiss Axioplan2 upright microscope. Images were taken with a 10X objective (NAD=D0.3), and micrographs acquired using a Zeiss AxioCam MRM CCD camera. Binary images were created in Fiji from manual tracings of the entire dendritic arbor for each neuron, and analyzed using the Fiji Sholl analysis plug- in. Sholl analysis was performed using 10-μm incremental increases in concentric circular diameter from the soma center. Only healthy neurons with homogenous GFP staining, without signs of dendritic blebbing and intact secondary and tertiary apical and basilar dendrites were imaged. Excitatory and inhibitory neurons were classified based on their distinctive morphologies. Excitatory neurons were identified by their prominent and highly branched apical dendrite, and numerous thinner and shorter basal dendrites covered by abundant dendritic spines. On the other hand, inhibitory neurons were recognized by having around 4-5 primary dendrites with similar thickness and length (multipolar), which further divide. All data were collected and analyzed by experimenters who were blind to experimental treatments.

### Expression and purification of sCNTNAP2

The construct encoding the entire extracellular domain of CNTNAP2 from residue 28 to 1261 was fused to the Fc portion of human IgG1 as detailed elsewhere (7, 59). This construct was transfected into HEK293GnTI- cells and the cells were selected by growth in the antibiotic G418. For large scale protein expression, stable cells were maintained at 37 °C and 5% (v/v) CO2 in Dulbecco’s modified Eagle’s medium containing up to 5% (v/v) FBS. CNTNAP2- Fc secreted in the medium was affinity purified using ProteinA-CaptivA TM PriMAB (RepliGen) and subsequently cleaved with recombinant 3C protease to remove the Fc fragment and leave the mature (i.e., no leader peptide) extracellular domain of CNTNAP2. The purified protein was buffer exchanged into 10 mM Hepes, pH 7.4, and 150 mM NaCl, concentrated to ∼3 to ∼6 mg/mL, and stored at 4 °C until used, as previously described (7).

### Coimmunoprecipitation assay

HEK293T cells were transfected with HA-MMP9 and Flag-CNTNAP2-WT or Flag-CNTNAP2 T1278I using Lipofectamine 2000 (ThermoFisher) in 100 mm dishes. After transfection, cells were treated with lysis buffer (50 mM Tris pH 7.5, 1% Triton X-100, 150 mM NaCl, 5 mM EDTA) containing protease inhibitor cocktail (ThermoFisher) and solubilized for 1 h at 4°C. Solubilized material was centrifuged at 10,000 g for 10 minutes at 4°C and the supernatants were incubated overnight at 4°C with 3 μg of anti-Flag rabbit antibody. Subsequently, 50 μl of slurry Sepharose A/G beads (SigmaAldrich) were added, and the mixture was incubated for 2 h at 4°C. Prior to use, A/G beads were washed and blocked for 1h with lysis buffer with 3% BSA. The complexes were washed three times with lysis buffer. Samples were then denatured for 5 min at 95°C with 50 μl of 2X sample buffer (BioRad, cat#1610737EDU) with β- mercaptoethanol. The immunoprecipitated proteins were separated using 4- 20% SDS-PAGE precast protein gels (BioRad) and blotted with anti-Flag and anti-HA antibodies.

### Quantification and statistical analysis

Statistical analysis was performed using GraphPad Prism. To compare differences between two groups, distributions of the measures were assessed for normality, and then Student’s *t*-test or Mann Whitney were used according to the distribution of the data. Data distribution was considered Gaussian after passing the Kolmogorov-Smirnov test. To compare multiple variables, a two-way ANOVA followed by multiple comparisons test was used. Statistical significance was defined as *P<*0.05. All statistical tests used were two-sided. Statistical details can be found in the figure legends.

## AVAILABILITY OF DATA AND MATERIALS

The datasets used and/or analyzed during the current study are available from the corresponding author on reasonable request.

## AUTHORS’ CONTRIBUTIONS

ML, LED, EM, NHP, PBS, SS, MDS, and MDMS conducted experiments; MDS, PP and MDMS designed experiments; ML, LED, EM, NP, PBS, SS, MDS, PP and MDMS analyzed the data; MDMS wrote the paper.

---

**Suppl. Fig. 1. Bioinformatic analysis comparing proteins across experimental groups. A.** Comparison of the sheddome between control and whisker trimming (Wtr) groups.. **B.** Comparison of the sheddome in the control vs. Wtr group in females only. **C.** Comparison of the sheddome in the control vs. Wtr group in males only. **D.** Comparison of the sheddome between female and male Wtr groups. **E.** Volcano plot showing differentially regulated proteins in the female Wtr vs. male Wtr sheddomes. **F.** Table of protein presence/absence: proteins unique to female Wtr (left, 29) and proteins unique to male Wtr (right, 34). **G.** GO (Gene Ontology) analysis of differentially regulated proteins identified in the volcano plot, including proteins unique to each group. Proteins detected in at least two of the three samples were considered “present” in the sheddome.

## REFERENCES

1. Malinverno M, Carta M, Epis R, Marcello E, Verpelli C, Cattabeni F, et al. Synaptic localization and activity of ADAM10 regulate excitatory synapses through N-cadherin cleavage. The Journal of neuroscience : the official journal of the Society for Neuroscience. 2010;30(48):16343–55.

2. Prox J, Bernreuther C, Altmeppen H, Grendel J, Glatzel M, D’Hooge R, et al. Postnatal disruption of the disintegrin/metalloproteinase ADAM10 in brain causes epileptic seizures, learning deficits, altered spine morphology, and defective synaptic functions. The Journal of neuroscience : the official journal of the Society for Neuroscience. 2013;33(32):12915–28, 28a.

3. Kuhn PH, Colombo AV, Schusser B, Dreymueller D, Wetzel S, Schepers U, et al. Systematic substrate identification indicates a central role for the metalloprotease ADAM10 in axon targeting and synapse function. Elife. 2016;5.

4. Peixoto RT, Kunz PA, Kwon H, Mabb AM, Sabatini BL, Philpot BD, et al. Transsynaptic signaling by activity-dependent cleavage of neuroligin-1. Neuron. 2012;76(2):396–409.

5. Suzuki K, Hayashi Y, Nakahara S, Kumazaki H, Prox J, Horiuchi K, et al. Activity-dependent proteolytic cleavage of neuroligin-1. Neuron. 2012;76(2):410–22.

6. Nagappan-Chettiar S, Johnson-Venkatesh EM, Umemori H. Activity-dependent proteolytic cleavage of cell adhesion molecules regulates excitatory synaptic development and function. Neurosci Res. 2017;116:60–9.

7. Martín-de-Saavedra MD, Dos Santos M, Culotta L, Varea O, Spielman BP, Parnell E, et al. Shed CNTNAP2 ectodomain is detectable in CSF and regulates Ca(2+) homeostasis and network synchrony via PMCA2/ATP2B2. Neuron. 2022;110(4):627–43.e9.

8. Lobete M, Salinas T, Izquierdo-Bermejo S, Socas S, Oset-Gasque MJ, Martin-de-Saavedra MD. A methodology to globally assess ectodomain shedding using soluble fractions from the mouse brain. Front Psychiatry. 2024;15:1367526.

9. Tushaus J, Muller SA, Kataka ES, Zaucha J, Sebastian Monasor L, Su M, et al. An optimized quantitative proteomics method establishes the cell type-resolved mouse brain secretome. EMBO J. 2020;39(20):e105693.

10. Kawai T, Elliott KJ, Scalia R, Eguchi S. Contribution of ADAM17 and related ADAMs in cardiovascular diseases. Cell Mol Life Sci. 2021.

11. Lichtenthaler SF, Lemberg MK, Fluhrer R. Proteolytic ectodomain shedding of membrane proteins in mammals-hardware, concepts, and recent developments. EMBO J. 2018;37(15).

12. Weber S, Saftig P. Ectodomain shedding and ADAMs in development. Development. 2012;139(20):3693–709.

13. Altmeppen HC, Prox J, Krasemann S, Puig B, Kruszewski K, Dohler F, et al. The sheddase ADAM10 is a potent modulator of prion disease. Elife. 2015;4.

14. Esch FS, Keim PS, Beattie EC, Blacher RW, Culwell AR, Oltersdorf T, et al. Cleavage of amyloid beta peptide during constitutive processing of its precursor. Science. 1990;248(4959):1122-4.

15. Vassar R, Bennett BD, Babu-Khan S, Kahn S, Mendiaz EA, Denis P, et al. Beta-secretase cleavage of Alzheimer’s amyloid precursor protein by the transmembrane aspartic protease BACE. Science. 1999;286(5440):735-41.

16. Schlepckow K, Kleinberger G, Fukumori A, Feederle R, Lichtenthaler SF, Steiner H, et al. An Alzheimer-associated TREM2 variant occurs at the ADAM cleavage site and affects shedding and phagocytic function. EMBO Mol Med. 2017;9(10):1356–65.

17. Vilalta A, Zhou Y, Sevalle J, Griffin JK, Satoh K, Allendorf DH, et al. Wild-type sTREM2 blocks Aβ aggregation and neurotoxicity, but the Alzheimer’s R47H mutant increases Aβ aggregation. J Biol Chem. 2021;296:100631.

18. Hampe W, Riedel IB, Lintzel J, Bader CO, Franke I, Schaller HC. Ectodomain shedding, translocation and synthesis of SorLA are stimulated by its ligand head activator. J Cell Sci. 2000;113 Pt 24:4475–85.

19. Fazeli E, Child DD, Bucks SA, Stovarsky M, Edwards G, Rose SE, et al. A familial missense variant in the Alzheimer’s disease gene SORL1 impairs its maturation and endosomal sorting. Acta Neuropathol. 2024;147(1):20.

20. Katz LC, Shatz CJ. Synaptic activity and the construction of cortical circuits. Science. 1996;274(5290):1133-8.

21. Fox K, Wong RO. A comparison of experience-dependent plasticity in the visual and somatosensory systems. Neuron. 2005;48(3):465–77.

22. Rochefort C, Gheusi G, Vincent JD, Lledo PM. Enriched odor exposure increases the number of newborn neurons in the adult olfactory bulb and improves odor memory. The Journal of neuroscience : the official journal of the Society for Neuroscience. 2002;22(7):2679–89.

23. Briner A, De Roo M, Dayer A, Muller D, Kiss JZ, Vutskits L. Bilateral whisker trimming during early postnatal life impairs dendritic spine development in the mouse somatosensory barrel cortex. J Comp Neurol. 2010;518(10):1711–23.

24. Zhang J, Cai F, Lu R, Xing X, Xu L, Wu K, et al. CNTNAP2 intracellular domain (CICD) generated by gamma-secretase cleavage improves autism-related behaviors. Signal Transduct Target Ther. 2023;8(1):219.

25. Poot M. Connecting the CNTNAP2 Networks with Neurodevelopmental Disorders. Molecular Syndromology. 2015;6(1):7–22.

26. Rodenas-Cuadrado P, Ho J, Vernes SC. Shining a light on CNTNAP2: complex functions to complex disorders. Eur J Hum Genet. 2014;22(2):171–8.

27. Rossi E, Verri AP, Patricelli MG, Destefani V, Ricca I, Vetro A, et al. A 12Mb deletion at 7q33-q35 associated with autism spectrum disorders and primary amenorrhea. Eur J Med Genet. 2008;51(6):631–8.

28. Poot M, Beyer V, Schwaab I, Damatova N, Van’t Slot R, Prothero J, et al. Disruption of CNTNAP2 and additional structural genome changes in a boy with speech delay and autism spectrum disorder. Neurogenetics. 2010;11(1):81–9.

29. Bakkaloglu B, O’Roak BJ, Louvi A, Gupta AR, Abelson JF, Morgan TM, et al. Molecular cytogenetic analysis and resequencing of contactin associated protein-like 2 in autism spectrum disorders. Am J Hum Genet. 2008;82(1):165–73.

30. O’Roak BJ, Deriziotis P, Lee C, Vives L, Schwartz JJ, Girirajan S, et al. Exome sequencing in sporadic autism spectrum disorders identifies severe de novo mutations. Nature Genetics. 2011;43(6):585–9.

31. Alarcón M, Abrahams BS, Stone JL, Duvall JA, Perederiy JV, Bomar JM, et al. Linkage, association, and gene-expression analyses identify CNTNAP2 as an autism-susceptibility gene. Am J Hum Genet. 2008;82(1):150–9.

32. Poliak S, Salomon D, Elhanany H, Sabanay H, Kiernan B, Pevny L, et al. Juxtaparanodal clustering of Shaker-like K+ channels in myelinated axons depends on Caspr2 and TAG-1. J Cell Biol. 2003;162(6):1149–60.

33. Anderson GR, Galfin T, Xu W, Aoto J, Malenka RC, Südhof TC. Candidate autism gene screen identifies critical role for cell-adhesion molecule CASPR2 in dendritic arborization and spine development. Proc Natl Acad Sci U S A. 2012;109(44):18120–5.

34. Gao R, Piguel NH, Melendez-Zaidi AE, Martin-de-Saavedra MD, Yoon S, Forrest MP, et al. CNTNAP2 stabilizes interneuron dendritic arbors through CASK. Molecular Psychiatry. 2018;23(9):1832–50.

35. St George-Hyslop F, Haneklaus M, Kivisild T, Livesey FJ. Loss of CNTNAP2 Alters Human Cortical Excitatory Neuron Differentiation and Neural Network Development. Biological Psychiatry. 2023;94(10):780–91.

36. Varea O, Martin-de-Saavedra MD, Kopeikina KJ, Schurmann B, Fleming HJ, Fawcett-Patel JM, et al. Synaptic abnormalities and cytoplasmic glutamate receptor aggregates in contactin associated protein-like 2/Caspr2 knockout neurons. Proc Natl Acad Sci U S A. 2015;112(19):6176–81.

37. Gdalyahu A, Lazaro M, Penagarikano O, Golshani P, Trachtenberg JT, Geschwind DH. The Autism Related Protein Contactin-Associated Protein-Like 2 (CNTNAP2) Stabilizes New Spines: An In Vivo Mouse Study. PLoS One. 2015;10(5):e0125633.

38. Fernandes D, Santos SD, Coutinho E, Whitt JL, Beltrão N, Rondão T, et al. Disrupted AMPA Receptor Function upon Genetic- or Antibody-Mediated Loss of Autism-Associated CASPR2. Cerebral Cortex. 2019;29(12):4919–31.

39. Martin-de-Saavedra MD, dos Santos M, Varea O, Spielman BP, Gao R, Forrest M, et al. CNTNAP2 ectodomain, detected in neuronal and CSF sheddomes, modulates Ca^2+^ dynamics and network synchrony. bioRxiv. 2019:605378.

40. Uhlhaas PJ, Singer W. Neural Synchrony in Brain Disorders: Relevance for Cognitive Dysfunctions and Pathophysiology. Neuron. 2006;52(1):155–68.

41. Uhlhaas PJ, Singer W. What Do Disturbances in Neural Synchrony Tell Us About Autism? Biological Psychiatry. 2007;62(3):190–1.

42. Uhlhaas PJ, Singer W. Abnormal neural oscillations and synchrony in schizophrenia. Nature Reviews Neuroscience. 2010;11(2):100–13.

43. Genovese G, Fromer M, Stahl EA, Ruderfer DM, Chambert K, Landén M, et al. Increased burden of ultra-rare protein-altering variants among 4,877 individuals with schizophrenia. Nature Neuroscience. 2016;19(11):1433–41.

44. Meta-analysis of GWAS of over 16,000 individuals with autism spectrum disorder highlights a novel locus at 10q24.32 and a significant overlap with schizophrenia. Mol Autism. 2017;8:21.

45. Zheng JJ, Li SJ, Zhang XD, Miao WY, Zhang D, Yao H, et al. Oxytocin mediates early experience-dependent cross-modal plasticity in the sensory cortices. Nat Neurosci. 2014;17(3):391–9.

46. Arakawa H, Erzurumlu RS. Role of whiskers in sensorimotor development of C57BL/6 mice. Behav Brain Res. 2015;287:146–55.

47. Strauss KA, Puffenberger EG, Huentelman MJ, Gottlieb S, Dobrin SE, Parod JM, et al. Recessive symptomatic focal epilepsy and mutant contactin-associated protein-like 2. N Engl J Med. 2006;354(13):1370–7.

48. Falivelli G, De Jaco A, Favaloro FL, Kim H, Wilson J, Dubi N, et al. Inherited genetic variants in autism-related CNTNAP2 show perturbed trafficking and ATF6 activation. Hum Mol Genet. 2012;21(21):4761–73.

49. Canali G, Garcia M, Hivert B, Pinatel D, Goullancourt A, Oguievetskaia K, et al. Genetic variants in autism-related CNTNAP2 impair axonal growth of cortical neurons. Hum Mol Genet. 2018;27(11):1941–54.

50. Hinkle CL, Diestel S, Lieberman J, Maness PF. Metalloprotease-induced ectodomain shedding of neural cell adhesion molecule (NCAM). J Neurobiol. 2006;66(12):1378–95.

51. Maretzky T, Schulte M, Ludwig A, Rose-John S, Blobel C, Hartmann D, et al. L1 is sequentially processed by two differently activated metalloproteases and presenilin/gamma-secretase and regulates neural cell adhesion, cell migration, and neurite outgrowth. Mol Cell Biol. 2005;25(20):9040–53.

52. Wierda KDB, Toft-Bertelsen TL, Gøtzsche CR, Pedersen E, Korshunova I, Nielsen J, et al. The soluble neurexin-1β ectodomain causes calcium influx and augments dendritic outgrowth and synaptic transmission. Scientific Reports. 2020;10(1):18041.

53. Naus S, Richter M, Wildeboer D, Moss M, Schachner M, Bartsch JW. Ectodomain shedding of the neural recognition molecule CHL1 by the metalloprotease-disintegrin ADAM8 promotes neurite outgrowth and suppresses neuronal cell death. J Biol Chem. 2004;279(16):16083–90.

54. Pischedda F, Piccoli G. The IgLON Family Member Negr1 Promotes Neuronal Arborization Acting as Soluble Factor via FGFR2. Front Mol Neurosci. 2015;8:89.

55. Gao R, Pratt CP, Yoon S, Martin-de-Saavedra MD, Forrest MP, Penzes P. CNTNAP2 is targeted to endosomes by the polarity protein PAR3. European Journal of Neuroscience. 2020;51(4):1074–86.

56. Huang da W, Sherman BT, Lempicki RA. Systematic and integrative analysis of large gene lists using DAVID bioinformatics resources. Nat Protoc. 2009;4(1):44–57.

57. Gao R, Piguel NH, Melendez-Zaidi AE, Martin-de-Saavedra MD, Yoon S, Forrest MP, et al. CNTNAP2 stabilizes interneuron dendritic arbors through CASK. Mol Psychiatry. 2018;23(9):1832–50.

58. Srivastava DP, Woolfrey KM, Penzes P. Analysis of dendritic spine morphology in cultured CNS neurons. Journal of visualized experiments : JoVE. 2011(53):e2794.

59. Rubio-Marrero EN, Vincelli G, Jeffries CM, Shaikh TR, Pakos IS, Ranaivoson FM, et al. Structural Characterization of the Extracellular Domain of CASPR2 and Insights into Its Association with the Novel Ligand Contactin1. J Biol Chem. 2016;291(11):5788–802.

