## Supplementary material for "ASD-Associated *CNTNAP2* Variants Disrupt Neuronal Arborization Through Impaired Regulation by Ectodomain Shedding": Suppl. Fig. 1

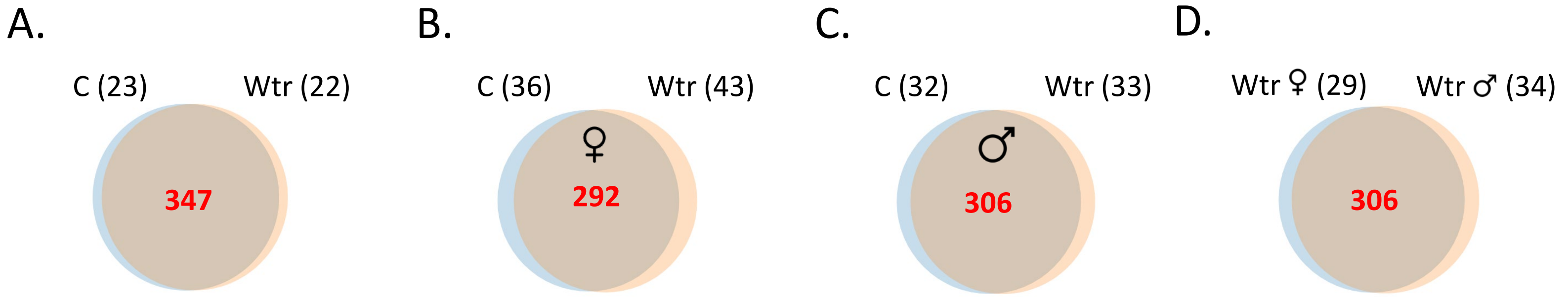


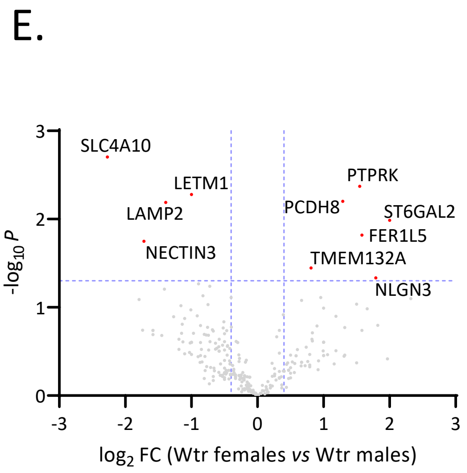


F.

| Wtr ♀ (29) | | Wtr ♂ (34) | |
|---|---|---|---|
| ABHD14A | FRAS1 | CD248 | SCARB2 |
| ANKLE2 | KCNN3 | CNTFR | SHISA9 |
| ANO8 | MRC2 | DAGLA | SLC1A3 |
| BTBD11 | PTPRE | ENPP6 | SLC25A30 |
| C2CD2L | REEP1 | GABBR2 | SLC39A12 |
| CD99 | SCN3B | GFRA2 | SLC6A11 |
| CHADL | SERINC3 | GJA1 | SLC7A6 |
| CLIC5 | SLC12A2 | GOSR1 | SLC8A1 |
| CLIC6 | SLC4A2 | HEG1 | SPRN |
| CLMN | SORCS1 | KCNB2 | SYT13 |
| CNTNAP5A | VDAC1 | LRFN2 | TENM4 |
| COQ8B | | LRTM2 | TMC5 |
| CUX1 | | MARCHF1 | UQCRFS1 |
| DCHS1 | | MEGF10 | VAMP4 |
| FAIM2 | | PCDHA10 | VSTM2B |
| FAM172A | | PLAAT3 | VTI1B |
| FLRT3 | | PLXNB3 | |
| FOCAD | | RETREG1 | |

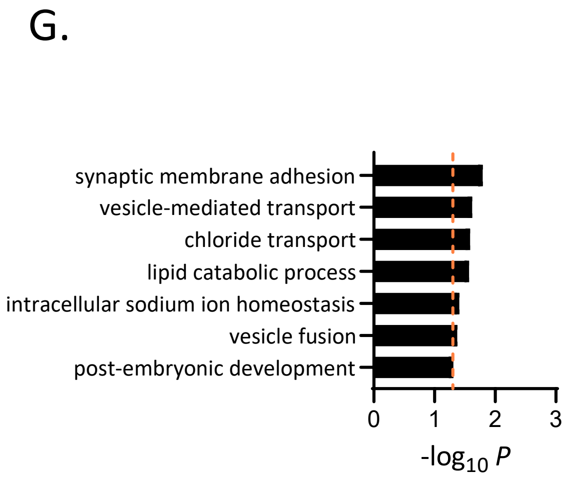
